# Population genomics of the maize pathogen *Ustilago maydis*: demographic history and role of virulence clusters in adaptation

**DOI:** 10.1101/2020.12.21.423782

**Authors:** Gabriel Schweizer, Muhammad Bilal Haider, Gustavo V. Barroso, Nicole Rössel, Karin Münch, Regine Kahmann, Julien Y. Dutheil

**Author notes:** Authors for correspondence: Gabriel Schweizer, University of Zürich, Department of Evolutionary Biology and Environmental Studies, Winterthurerstrasse 190, 8057 Zürich, Switzerland, Julien Y. Dutheil, Max-Planck-Institute for Evolutionary Biology, Research Group Molecular Systems Evolution, August-Thienemann-Straße 2, 24306 Plön, Germany.

## Abstract

The tight interaction between pathogens and their hosts results in reciprocal selective forces that impact the genetic diversity of the interacting species. The footprints of this selection differ between pathosystems because of distinct life-history traits, demographic histories, or genome architectures. Here, we studied the genome-wide patterns of genetic diversity of 22 isolates of the causative agent of the corn smut disease, *Ustilago maydis*, originating from five locations in Mexico, the presumed center of origin of this species. In this species, many genes encoding secreted effector proteins reside in so-called virulence clusters in the genome, an arrangement that is so far not found in other filamentous plant pathogens. Using a combination of population genomic statistical analyses, we assessed the geographical, historical and genome-wide variation of genetic diversity in this fungal pathogen.

We report evidence of two partially admixed subpopulations that are only loosely associated with geographic origin. Using the multiple sequentially Markov coalescent model, we inferred the demographic history of the two pathogen subpopulations over the last 0.5 million years. We show that both populations experienced a recent strong bottleneck starting around 10,000 years ago, coinciding with the assumed time of maize domestication. While the genome average genetic diversity is low compared to other fungal pathogens, we estimated that the rate of non-synonymous adaptive substitutions is three times higher in genes located within virulence clusters compared to non-clustered genes, including non-clustered effector genes. These results highlight the role that these singular genomic regions play in the evolution of this pathogen.

**Significance statement:** The maize pathogen *Ustilago maydis* is a model species to study fungal cell biology and biotrophic host-pathogen interactions. Population genetic studies of this species, however, were so far restricted to using a few molecular markers, and genome-wide comparisons involved species that diverged more than 20 million years ago. Here, we sequenced the genomes of 22 Mexican *U. maydis* isolates to study the recent evolutionary history of this species. We identified two co-existing populations that went through a recent bottleneck and whose divergence date overlaps with the time of maize domestication. Contrasting the patterns of genetic diversity in different categories of genes, we further showed that effector genes in virulence clusters display a high rate of adaptive mutations, highlighting the importance of these effector arrangements for the adaptation of *U. maydis* to its host.

## Introduction

The coevolution between plant pathogens and their hosts impacts the genetic diversity of the interacting species. The response to these reciprocal selective forces depends on multiple factors, including the genome architecture of the organisms (e.g. genome size and karyotype structure), their life history traits (e.g. importance and frequency of sexual reproduction), but also stochastic factors in relation to demography (e.g. variable population size and population structure). Consequently, plant domestication and the subsequent emergence of agriculture, which typically results in a population bottleneck and a strong directional selection, had a strong impact on the selected organisms (Tang *et al*., 2010; Milla *et al*., 2015). It also affected the evolution of the associated pathogens, because domestication resulted in significant losses of genetic variation and strong selection on a few genes (Glemin and Bataillon, 2009). For example, speciation of the rice blast pathogen *Magnaporthe oryzae*, the wheat pathogen *Zymoseptoria tritici* and the barley pathogen *Rhynchosporium secalis* from their wild relatives was associated with the domestication of their host plants (Couch *et al*., 2005; Stukenbrock *et al*., 2007; Zaffarano *et al*., 2008). Here, we investigate the evolutionary history of the maize pathogen *Ustilago maydis*, a basidiomycete from the group of smut fungi (family: Ustilaginaceae). This family comprises about 550 described species (Begerow *et al*. 2014), among which are pathogenic species of grasses, including crops like maize, sorghum, wheat, barley, and sugarcane (Agrios, 2005). The genomes of several crop pathogens have been sequenced (Kämper *et al*., 2006; Schirawski *et al*., 2010; Laurie *et al*., 2012; Que *et al*., 2014; Taniguti *et al*., 2015; Dutheil *et al*., 2016; Benevenuto *et al*., 2018) as well as some species parasitizing wild grasses or dicot plants (Sharma *et al*., 2014; Rabe *et al*., 2016; Ye *et al*., 2017). Dating of speciation events between these species suggested that their divergence predates the domestication of their hosts and therefore occurred in their wild ancestors (Munkacsi *et al*., 2007; Schweizer *et al*., 2018). Among these species, *U. maydis* is the best studied and serves as a model for elucidating the molecular basis of biotrophic host-pathogen interactions (Matei and Doehlemann, 2016; Lanver *et al*., 2017). These studies showed that the interaction with the host plant maize is largely controlled by secreted effector proteins of which about half lack known functional domains (Lanver *et al*., 2017). Genome comparisons of *U. maydis* and related species revealed that many effector genes reside in gene clusters in the genome, an arrangement that is so far not described in other filamentous plant pathogens. Despite the high divergence level of these clustered effectors, homology between clusters of distinct species was established due to conserved synteny between genomes (Schirawski *et al*., 2010; Dutheil *et al*., 2016). Functional analyses of such clusters showed that they contain important virulence determinants in the barley pathogen *Ustilago hordei* (Ali *et al*., 2014), in *U. maydis* (Kämper *et al*., 2006; Schirawski *et al*., 2010; Brefort *et al*., 2014; Navarrete *et al*., 2019), and in the maize pathogen *Sporisorium reilianum* (Ghareeb *et al*., 2018).

It is hypothesized that the center of origin of *U. maydis* lies in Mexico from where it spread following the domestication of maize from teosinte (Munkacsi *et al*., 2008), starting 6,000 to 10,000 years ago (Matsuoka *et al*., 2002; Hake and Ross-Ibarra, 2015). Investigating infected maize fields demonstrated that *U. maydis* shows a single generation and limited spreading between host plants in one growing season (Baumgarten *et al*., 2007). Moreover, an analysis of amplified fragment length polymorphism markers of isolates sampled in the USA and Uruguay showed that *U. maydis* reproduces predominantly by out-crossing, and this finding was independent of differences in agricultural practice at the sampling sites (Barnes *et al*., 2004).

Munkacsi et al. (2008) investigated the impact of maize domestication on the evolution of *U. maydis*, using ten microsatellite markers. Samples from different locations revealed that subpopulations in Mexico diverged within a time window that is consistent with the domestication and cultivation of maize in the Americas. Moreover, genetic diversity of *U. maydis* was not found to be greater in Mexico (the presumed origin of the species) than in other parts of the Americas, suggesting that the domestication of maize from teosinte imposed a bottleneck that reduced the ancient genetic diversity in *U. maydis* (Munkacsi *et al*., 2008). Furthermore, analyses of sequence polymorphisms in 18 *U. maydis* isolates originating from 11 locations in Europe, North America, and South America with a focus on the virulence clusters 2A and 19A as well as the single effector *pep1* demonstrated low genetic variation in these regions and uncovered three subpopulations based on geographic origin (Kellner *et al*., 2014). While these studies of individual genomic loci highlighted the effect of domestication on the evolutionary history of *U. maydis*, they did not allow the detailed inference of the demographic history and genome-wide patterns of selection in this species. Such studies require the availability of full genome sequences (Stukenbrock *et al*., 2011; Grünwald *et al*., 2016).

To extend our understanding of the evolutionary history of *U. maydis*, we employed a population genomics approach and sequenced 22 isolates originating from five different regions in Mexico (Valverde *et al*., 2000). We used this data set to investigate the population structure and the demographic history of the sampled isolates. We further assessed patterns of genome-wide nucleotide diversity and inferred the rate of adaptive substitutions in distinct categories of genes, allowing us to highlight the unique role of virulence clusters in the adaptive evolution of this fungal plant pathogen.

## Materials and Methods

### Origin, genomic DNA extraction, and sequencing of haploid U. maydis isolates

We sequenced the genome of 22 Mexican *U. maydis* isolates that are part of an earlier described isolate collection (Valverde *et al*., 2000; supplementary table S1). The isolates were stored as haploid sporidia in glycerol stocks at −80°C. Isolates were thawed by plating them on Potato-Dextrose (PD) plates (3.9% [w/v] Potato-Dextrose Agar, 1% [v/v] Tris–HCl [1 M, pH 8.0]) and incubating them for two days at 28°C. Next, fungal cells were scratched off the PD plates and ground together with glass beads in liquid nitrogen. Genomic DNA was extracted by adding 500 µl TE-phenol/chloroform (1:1) and 500 µl lysis buffer (100 mM NaCl, 10 mM Tris-HCl (pH 8.0), 1 mM EDTA, 2 % Triton X-100, and 1 % SDS) followed by precipitation in 70% Ethanol. RNA was removed from the samples with the Master Pure Complete DNA & RNA Purification Kit (Biozym Scientific, Hessisch Oldendorf, Germany). DNA concentration was adjusted to about 150 to 400 ng/µl and about 1 µg of DNA was used for sequencing. After fragmentation of the genomic DNA, sequencing libraries were prepared using the TruSeq DNA LT Kit (Illumina, San Diego, USA) and sequenced at the Max Planck Genome Centre (Cologne, Germany) using the HiSeq sequencing kit on a HiSeq2000 cycler (Illumina). Paired-end sequencing was performed with a 100 base pair read length and for each sequencing library, at least 21.6 million reads were generated, corresponding to a 100-fold average coverage (supplementary table S1). All Illumina paired-end reads were deposited at NCBI (BioProject ID: PRJNA561077).

### De novo genome assemblies

A d*e novo* assembly of the haploid genome was generated individually for each Mexican isolate with SOAPdenovo2 (Luo *et al*., 2012) as follows: all odd kmer lengths ranging from 51 to 83 were tested and the kmer length yielding the highest N_50_ was selected for each library. The estimated genome size (option -z) was set in all cases to 20,000,000 base pairs, the genome size of the reference genome for *U. maydis* isolate 521 (Kämper *et al*., 2006). Resulting genome assembly statistics, including N_50_ contig lengths for each kmer and isolate are summarized in supplementary table S2. Next, SOAPdenovo2 was used to get a genome for each isolate with the determined optimal kmer length. A sparse pregraph was built, and contigs were then computed, mapped and assembled into scaffolds. Finally, the GapCloser program was used to close remaining assembly gaps. The assembled genome sequences were deposited at NCBI (BioProject ID: PRJNA561077).

### Multiple genome alignment

We generated a multiple genome alignment that comprised the *de novo* assembled genomes of the 22 Mexican *U. maydis* isolates together with the *U. maydis* reference genome of the isolate 521 (Kämper *et al*., 2006). The reference genome sequence was obtained from Mycocosm of the Joint Genome Institute (Grigoriev *et al*., 2014) in version 2_2 from December 5, 2017. The 23 genomes served as input for the Multiz genome aligner from the Threaded Blockset Aligner package (Blanchette *et al*., 2004). The resulting alignment was then projected on the reference genome, yielding an alignment length of 20,028,090 base pairs. This alignment was then processed using MafFilter (Dutheil *et al*., 2014). First, all synteny blocks were realigned using Mafft (Katoh and Standley, 2013), with blocks of a length greater than 10 kb being first split before alignment for computational efficiency. This unfiltered alignment was then subjected to two pipelines. The first pipeline focused on protein coding genes and extracted all exons from the unfiltered alignment (see below, *Building gene families*). In the second pipeline, the realigned synteny blocks were filtered to remove ambiguously aligned regions. This was achieved in two steps: first, only blocks that comprised sequences from all 23 isolates are kept and alignment blocks with multiple “paralogous” sequences per species were discarded. Second, alignment blocks were further processed with a sliding window approach. Within 10 bp windows slid by 1 nucleotide, short indels were identified and the window was discarded if it contained at least one indel shared by at least two isolates, or, alternatively, if the quantity of gap characters in the 23 species was higher than 100 gaps in the window. In these two steps, unresolved base positions were assigned as gaps. These filtering steps yielded a final alignment with a length of 19,224,664 bp in 2,676 blocks. Alignment lengths and number of blocks resulting from each filtering step are summarized in supplementary table S3. We generated a second alignment using the same protocol, including this time all 23 *U. maydis* sequences together with the genome sequence of *S. reilianum* SRZ2 (version 2) which we obtained from the PEDANT database (Walter *et al*., 2009). The corresponding alignment statistics for each filtering step are provided in supplementary table S3. Pairwise similarity distances were computed using MafFilter and a global tree was constructed using the FastME software (Lefort *et al*., 2015), with default nucleotide model, nearest neighbor interchange (NNI) and subtree pruning regrafting (SPR) topology optimization. One thousand bootstrap replicates were performed in order to assess the support of each clade.

### Analyses of population structure

Single nucleotides polymorphisms (SNPs) were called from the filtered alignment using MafFilter and exported to a file in the Variant Call Format (VCF). The set of SNPs was thinned according to linkage disequilibrium using the bcftools (Li, 2011), and only pairs with r^2^ < 0.6 in 250 kb windows were kept. The resulting “unlinked” SNP set was exported to a file in PLINK format using plink 1.9 (Purcell *et al*., 2007). The smartPCA software (Patterson *et al*., 2006) was used to compute principal components from the unlinked SNP set, and results were plotted with the R statistical environment (Ortutay and Ortutay, 2017). Model-based inference of population structure was conducted using the ADMIXTURE software (Alexander *et al*., 2009) on SNPs filtered as for the PCA analysis. We performed a cross-validation analysis for a range of models with one to six genetic components. Each model was rerun from ten random initial conditions and results were summarized using the PONG software (Behr *et al*., 2016).

### Analyses of nucleotide diversity

The mean number of nucleotide difference between all pairs of sequences (π), GC content and fixation index F_ST_ were computed in non-overlapping windows of 10 kb from the multiple genome alignment of *U. maydis* isolates. The divergence between *U. maydis* and *S. reilianum* reference genomes was computed from the multiple genome alignment with outgroup in non-overlapping windows of 10 kb. A Tamura 92 model (Tamura, 1992) was fitted independently in each window in order to account for multiple substitutions while accounting for variable ratios of transitions over transversions, as well as non-homogeneous GC content. F_ST_ values were calculated using Hudson’s 1992 estimator (Hudson *et al*., 1992). All calculations were performed using the MafFilter program (Dutheil *et al*., 2014). The distribution of F_ST_ values showed a tail of extreme F_ST_ values (supplementary figure S1) and was best fitted with a mixture of normal distributions using the ‘fitdistr’ function from the MASS package (Venables and Ripley, 2002) for R (https://www.R-project.org). To assess the significance of high F_ST_ values, we computed the probability that the F_ST_ value belonged to the lower mode of the distribution, using the estimated parameters of the two normal distributions. The 191 regions for which this probability was lower than 1% were considered as high F_ST_ regions and were subsequently scanned for genes, resulting in 751 candidate genes.

### Detection of Gene Ontology term enrichments

All annotated *U. maydis* proteins were used as input for an Interpro search with version 5.35-74.0-64 (Mitchell *et al*., 2019), and mapped Interpro domains for each protein are listed in supplementary table S4. Next, the Interpro domains were linked to Gene Ontology (GO) Terms with the file “interpro2go”, which is provided by the GO consortium (version 2019/05/02 15:27:19; http://current.geneontology.org/ontology/external2go/interpro2go). In this way, 1,948 unique GO terms could be assigned to 4,147 *U. maydis* proteins (supplementary table S4). Each GO Term was associated with one of the three major subontologies “Cellular Component,” “Biological Process,” or “Molecular Function” with the Bioconductor package topGO (Alexa *et al*., 2006). Enriched GO Terms were then identified by computing *P-*values for each GO term using Fisher’s classic test with parent-child correction (Grossmann *et al*., 2007). For this analysis, genes with a high F_ST_ value were compared to all genes for which F_ST_ values could be computed, and results were considered to be significant at the 5% level.

### Inference of demography using the multiple sequentially Markov coalescent

We used MSMC2, a re-implementation of the multiple sequentially Markov coalescent (Malaspinas *et al*., 2016) to estimate the time variation of coalescence rates. The filtered alignment was converted to MSMC input format using the MafFilter program (Dutheil *et al*., 2014). MSMC2 was then run with default options. In order to convert time estimates from coalescent units to years and coalescence rates into effective population sizes, measures of generation time and average mutation rate are needed. Munkacsi and colleagues provided estimates of the synonymous mutation rate in smut fungi using six species comparisons, in four different genes, leading to 24 estimates {x_i_} of the mutation rate (table 4 in Munkacsi *et al*., 2007). These estimates are exponentially distributed, and we therefore computed a genome geometric mean (*u*) using the formula *u* = exp(∑_*i*_ log(x_*i*_)/24), leading to a value of *u* = 5.23×10^−9^ mutations per site per generation. We further considered a generation time of one year (Munkacsi *et al*., 2008). Cross-coalescence rate analyses were conducted following the protocol described in the documentation of the MSMC2 program. In order to assess the significance of the pattern of cross-coalescence rate between two samples, we randomly permuted genomes in samples and recomputed the rates of cross-coalescence. A total of 10 permutations was conducted.

### Building gene families

We used the multiple genome alignment of *U. maydis* isolates to extract nucleotide sequences of protein coding genes according to the annotation of the *U. maydis* reference genome, obtained from Mycocosm of the Joint Genome Institute (Grigoriev *et al*., 2014). This annotation encompassed 6,785 protein coding genes, of which we discarded 36 genes with splice variants. We extracted nucleotide sequences for 6,742 genes (supplementary table S4). An outgroup sequence from the related species *Sporisorium reilianum* f. sp. *zeae* was further added using the procedure described below. We obtained the proteome of *S. reilianum* (Schirawski *et al*., 2010) from the protein data base PEDANT (Walter *et al*., 2009) with 6,676 proteins. The proteome of the *U. maydis* isolate 521 was searched against the *S. reilianum* proteome using blastp, and the result was used as input for the SiLiX algorithm in order to reconstruct gene families (Miele *et al*., 2011). This software infers homologous relationships based on two criteria: the percent identity between two sequences and the coverage, defined as the relative length of a hit compared to the total length of the two sequences. We used a range for coverage and identity thresholds between 5% and 95% in 5% steps to identify values that result in the largest number of families with one-to-one homologs. The thresholds of 45 % identity and 55 % coverage were selected, because they lead to the maximum number (5,685) of families comprising one gene each in *U. maydis* and *S. reilianum* (supplementary table S5). This allowed us to map a single *S. reilianum* ortholog to 5,678 genes that were extracted from the multiple genome alignment. Several proteins in virulence clusters, however, could not be assigned to families of 1:1 orthologs predicted by SiLiX, because such genes evolved by duplication (Dutheil *et al*., 2016). In order to identify an outgroup sequence for these genes, we conducted a second blastp search with, as query, all *U. maydis* genes that could be extracted from the multiple genome alignment but had not been mapped to a *S. reilianum* ortholog. We used all *S. reilianum* proteins that are not mapped to a *U. maydis* ortholog by SiLiX as target. We considered only hits with an E-value < 10^−6^ for further analyses and found 404 cases where one *U. maydis* gene mapped to one *S. reilianum* gene and vice versa. In summary, we could assign a *S. reilianum* outgroup sequence to 6,082 *U. maydis* genes out of 6,742 genes that could be extracted from the multiple genome alignment (supplementary table S4). This set of gene families was aligned at the codon level using MACSE (Ranwez *et al*., 2011), with the exception of one gene (*UMAG_10543*) for which the program failed to output an alignment. Two genes (*UMAG_00001* and *UMAG_10807*) were additionally discarded as their annotation changed since the previous version of the genome used in Dutheil *et* al. (2016) and they were, therefore, not included in the prediction of effector clusters. 6,051 genes were detected in all 22 Mexican isolates and were selected for subsequent analyses. Of these genes, 56 have at least one predicted in-frame stop codon before the end of the coding sequence in the reference genome and were discarded, as they could correspond to polymorphic open reading frames or contain sequencing errors. The final data set contained a total of 5,993 genes. Annotations of candidate effector genes were taken from the “strict” prediction described in Dutheil et al. (2016), which contained 553 genes predicted to encode secreted effector proteins. In the same study, 156 effector genes were found in gene clusters. The filtered dataset studied here contained 95 genes in virulence clusters, 344 non-clustered effectors, and 5,554 other non-effectors, non-clustered genes.

### Reconstruction of site frequency spectra

Filtered alignments were analyzed with the bppPopStat program from the bppSuite software (Guéguen *et al*., 2013) in order to compute the unfolded synonymous and non-synonymous site frequency spectra (SFS) for each gene of the 22 Mexican isolates. Ancestral alleles were inferred using a marginal reconstruction after fitting a codon model (Yang and Nielsen’s model with F3×4 frequencies; Yang and Nielsen, 1998), including the outgroup sequence. When computing the site frequency spectra, positions with more than two alleles were ignored, as well as positions where the outgroup displayed an allele distinct from the set of alleles in the ingroup.

### Estimation of the distribution of fitness effects (DFE) of mutations and rate of adaptive substitutions

Gene-specific SFS for each gene category (gene in virulence cluster, non-clustered effectors, or non-clustered, non-effector genes) set were obtained by pooling the SFS for each constitutive gene to generate the input file for the Grapes program (Galtier, 2016). A model selection procedure was conducted for each set separately, using the ‘-m all’ option of Grapes. We further used the ‘no_div_param’ option to use divergence predictions from the polymorphism data instead of using the sequence of the outgroup to estimate the corresponding parameters, relaxing the assumption that the DFE was constant since the divergence with the outgroup. The best model according to the Akaike’s information criterion (AIC) was found to be the scaled-beta distribution for the non-clustered effector genes, while the gamma-exponential distribution and gamma-gamma distributions best fitted clustered genes and non-effector non-clustered genes, respectively (see supplementary table S6). In all categories of genes, however, these three models had very similar AIC values, and we performed a model averaging procedure by weighting each model with its relative likelihood (Dormann *et al*., 2018). In order to assess the sampling variance of the inferred DFE and rate of adaptive substitutions for each set of genes, a bootstrap procedure was conducted by re-sampling genes in each category 100 times. Parameters of the DFE as well as rates of adaptive substitutions were estimated for each bootstrap replicate, using the model with the best fit for each gene category, and their distribution used to compute confidence intervals. To assess the significance of observed differences in parameters between gene categories, we performed a permutation test, re-sampling all genes between the three categories a 1,000 times and rerunning Grapes with the selected model on each sample. For each pair of gene sets, *P*-values were then computed using the formula (∑_*i*_[S_*i*_ ≥ S_*obs*_]+1)/1001 where S_*i*_ denotes the difference in parameter estimates for replicate *i* and ∑_*i*_[S_*i*_ ≥ S_*obs*_] is the number of replicates for which the difference is greater or equal to the observed difference. The following comparisons were performed: clustered genes *vs*. non-clustered effectors, clustered genes *vs*. non-clustered non-effector genes, and non-clustered effectors *vs*. non-clustered non-effector genes. The parameters tested included non-synonymous diversity (π_N_), synonymous diversity (π_S_), and their ratio (π_N_/π_S_), ratio of non-synonymous to synonymous divergences (d_N_/d_S_ = ω), the rate of non-adaptive non-synonymous substitutions (ω_NA_), the rate of adaptive non-synonymous substitutions (ω_A_), and the proportion of adaptive non-synonymous substitutions (α). In order to control for protein length, a similar analysis was conducted after selecting a subset of non-clustered effector genes and non-clustered, non-effector genes with protein length similar to that of clustered genes. This was achieved by selecting, for each clustered gene, the corresponding gene with the most similar protein length in the non-clustered effector (resp. non-clustered, non-effector) gene sets. The generated sets have, therefore, the same number of genes as the clustered set, and their average length did not differ significantly (Kruskal-Wallis rank sum test, *P*-value = 1, supplementary file S1).

### Analysis of mating type loci

Sequences from the *a* and *b* mating type loci were extracted from the genome alignment using annotations from the *U. maydis* reference genome. The corresponding genes are *UMAG_02382* (1 exon), *UMAG_02383* (4 exons), and UMAG_02384 (3 exons) for the *a*-locus on chromosome 5, and *UMAG_12052* (2 exons) and *UMAG_00578* (2 exons) for the *b*-locus on chromosome 1. The complete region was extracted for each locus. Sequences of the two genes of the *b*-locus were combined with publicly available sequences (Kämper *et al*., 2020), and phylogenetic trees were reconstructed for each gene using the PhyML program (Guindon *et al*., 2010), using a Le and Gascuel model of protein evolution (Le and Gascuel, 2008) with a 4-classes discrete Gamma distribution of rates. The “Best of NNI and SPR” topology search option was selected, and 100 non-parametric bootstraps were generated. Nodes with a bootstrap value below 60% were set as unresolved. Trees were plotted using the ‘ggtree’ package for R (Yu *et al*., 2016).

### Verification of the *a* mating type locus of isolate A

All reads were mapped on the assembly of the isolate A using bwa mem (Li and Durbin, 2009). Site-specific coverage and read heterogeneity were computed using the mpileup program from the samtools (Li *et al*., 2009) and further processed by python and R scripts. We performed a second independent genomic DNA extraction for isolates A and B. Isolates were streaked out from a glycerol stock and were cultivated for two days on PD plates at 28°C. Next, a single colony was inoculated in YEPS-light liquid medium (1 % w/v yeast extract, 1 % w/v peptone, 1 % w/v sucrose) and grown over night at 28°C. Cells were pelleted and lysed with ca. 0.3 g glass beads, 500 µl TE-phenol/chloroform (1:1) and 500 µl lysis buffer for 15 minutes on a Vibrax shaker. The supernatant was precipitated with 70 % (v/v) ethanol and dissolved in TE-buffer (1 mM Na_2_-EDTA, 10 mM Tris-HCl with pH 8.0) supplemented with RNaseA.

The Phusion high fidelity PCR master mix (New England BioLabs) was prepared with 0.2µM primers (forward sequence: CTAGCTACACCAGCGAGGACGATA; reverse sequence: TCATTCCTAGCTCTTCTTGCGTTGA) and 100 ng genomic DNA as template in a 20 µl reaction volume. The PCR was performed as follows: initial denaturation at 98°C for 1 minute; 30 cycles with10 seconds denaturation at 98°C, 15 seconds annealing at 67°C, 5 minutes extension at 72°C and 7 minutes final extension at 72°C. Products from the PCR were analyzed by gel electrophoresis. The entire volume of the samples was mixed with 3 µl of bromophenol blue dye and loaded on 1% agarose gel prepared in 1X TAE buffer (40 mM Tris, 20 mM acetic acid, 1 mM EDTA with pH 8.0).

## Results and Discussion

We sequenced 22 haploid isolates of *U. maydis* originating from five locations in Mexico (Valverde *et al*., 2000; figure 1A and supplementary table S1). We performed a *de novo* genome assembly for each isolate and computed a multiple genome alignment including the previously sequenced reference isolate 521 (Kämper *et al*., 2006). After filtering for alignment uncertainty, the length of the alignment totalized 19.2 Mb, covering 97.76 % of the *U. maydis* reference genome. A total of 61,745 SNPs was called from the newly sequenced isolates, corresponding to a mean number of nucleotide differences of 9×10^−4^ per nucleotide. This level of diversity is comparable to that found in populations of great apes or of *Drosophila melanogaster* (Nam *et al*., 2015; Haudry *et al*., 2020), but remarkably low for a fungal pathogen (Zheng *et al*., 2013; McMullan *et al*., 2018; Stukenbrock and Dutheil, 2018). Moreover, 6,742 genes out of the 6,785 annotated genes in the reference genome had a homologous sequence in all Mexican isolates, and of these, 5,993 had both a coding sequence without predicted in-frame stop codon and an identifiable homolog in *S. reilianum*. This high level of sequence conservation shows (1) that the generated dataset is of high quality and encompasses a large proportion of the genome of the pathogen and (2) that the genetic diversity of the population at the center of origin is very low.

**Figure 1:**
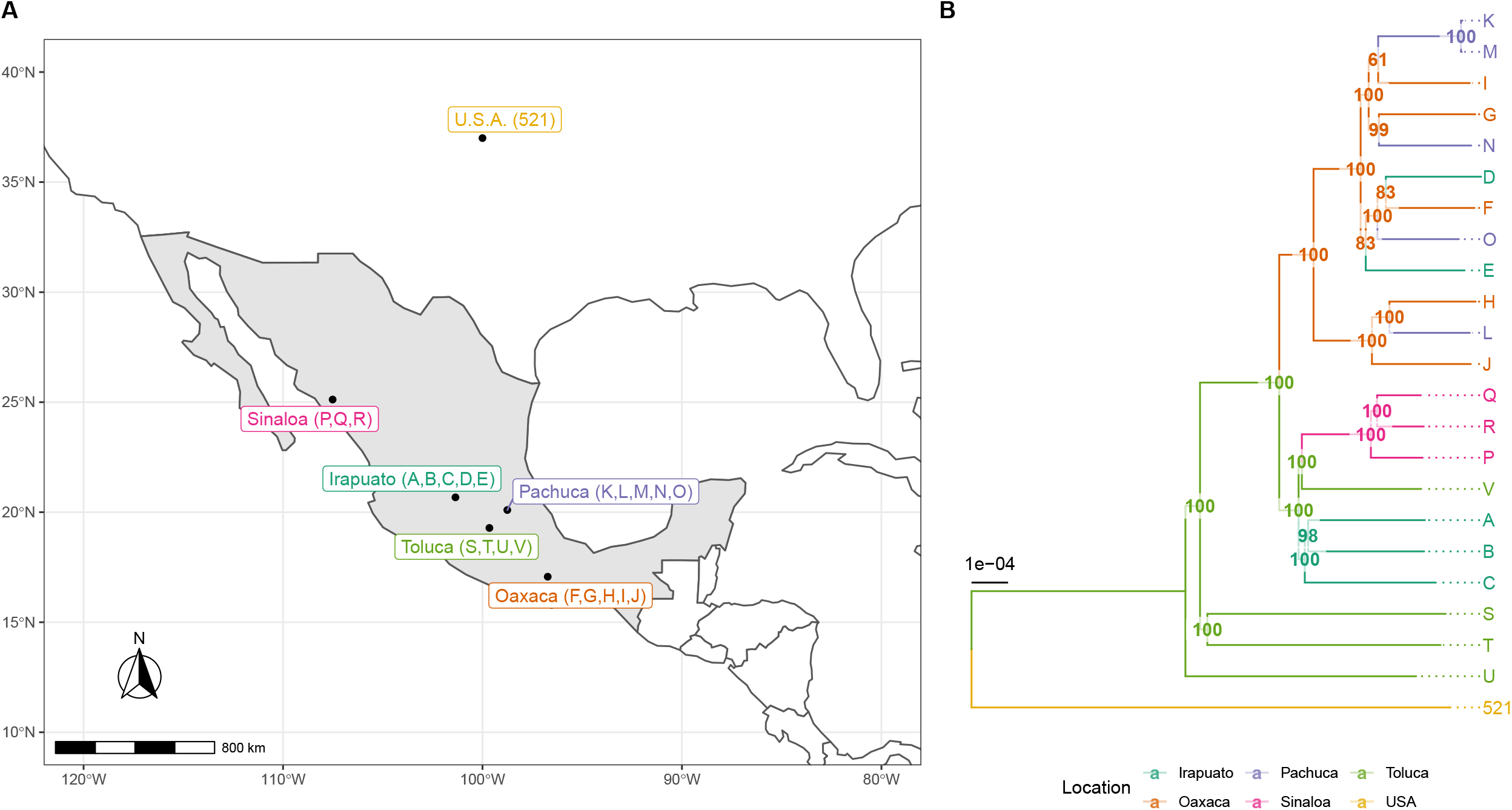
Genome similarity of isolates is only loosely associated with their geographic origin. (A) Mexican *U. maydis* isolates (named with letters from A to V) originated from the five regions Irapuato, Oaxaca, Pachuca, Sinaloa, and Toluca. The reference isolate 521 was collected in the USA. (B) Tree showing the genome-wide similarity levels between isolates. Isolates and branches are colored according to the sampling location as shown in A. Numbers indicate bootstrap support values as percentage from 1,000 replicates, and branch lengths are the mean number of substitutions per site.

### The investigated isolates represent two subpopulations

To infer the level of similarity between the sequenced isolates, we constructed a global tree (figure 1B). We found that all Mexican isolates (Valverde *et al*., 2000) are more similar to each other than they are to the genome of the reference isolate 521, which was collected from a corn field near St. Paul/Minnesota, USA (Holliday, 1961). Within the Mexican isolates, isolates P, Q, and R were all collected from Sinaloa and cluster in one group. Moreover, the isolates S, T, and U from Toluca clustered independently of all other isolates. Isolates M and K appeared to be very similar, displaying less than one SNP per chromosome on average after quality filtering, suggesting that the two individuals are very closely related. Overall, we detected only a loose association between sampling origin and genome similarity, suggesting that population structure, if any, is not induced by geography. To further investigate the population structure, we performed a principal component analysis (PCA) (Patterson *et al*., 2006). The results of the PCA were consistent with those of the global genome similarity tree (figure 2A and figure 2B): the first principal components distinguished three groups of isolates. The first set, consisting of isolates D, E, F, G, I, K, M, N, and O (referred below as the ‘DEFGIKMNO’ or ‘DEFGIKNO’ population, depending on whether the M isolate was included in the analysis, see below) formed a well-supported group in the similarity tree containing four of the five isolates from Pachuca, three of the five isolates from Oaxaca, and two of the five isolates from Irapuato. The second set, consisting of isolates A, B, C, P, Q, R, S, T, U, and V (referred below as the ‘ABCPQRSTUV’ population), grouped the four isolates from Toluca, the three isolates from Sinaloa, and three of the five isolates from Irapuato. The third group was found in between these two populations and consisted of only three isolates: isolates J and H from Oaxaca and isolate L from Pachuca. To corroborate this finding, we employed the ADMIXTURE program (Alexander *et al*., 2009). Cross-validation favored a model with two subpopulations. Rerunning the model estimation procedure with distinct initial conditions showed that this model is also the most consistent, that is, 10 replicates out of 10 agreed on the population partitioning (supplementary figure S2). The two inferred subpopulations matched the grouping of the PCA analysis and the similarity tree, showing that the isolates H, L and J are a mixture of these two subpopulations with a ratio of roughly 70% to 30% (figure 2C). Our findings are in line with previous studies that also reported the presence of subpopulations of *U. maydis* in Mexico (Munkacsi *et al*., 2008). Interestingly, the reference isolate 521 grouped together with the ABCPQRSTUV population, while showing some low degree of admixture with the DEFGIKMNO group. Deep sampling outside of Mexico is required to confirm the relationship of the reference isolate to the Mexican populations. The mechanisms of divergence of the two populations remain to be elucidated. Teosinte occurs in Mexico with two subspecies, *Zea mays* ssp. *parviglumis* and *Zea mays* ssp. *mexicana* (Fukunaga *et al*., 2005; Ross-Ibarra *et al*., 2009). An interesting hypothesis is that the two populations of *U. maydis*, while being able to infect maize, could represent formae speciales primarily adapted to each of these two subspecies of teosinte.

**Figure 2:**
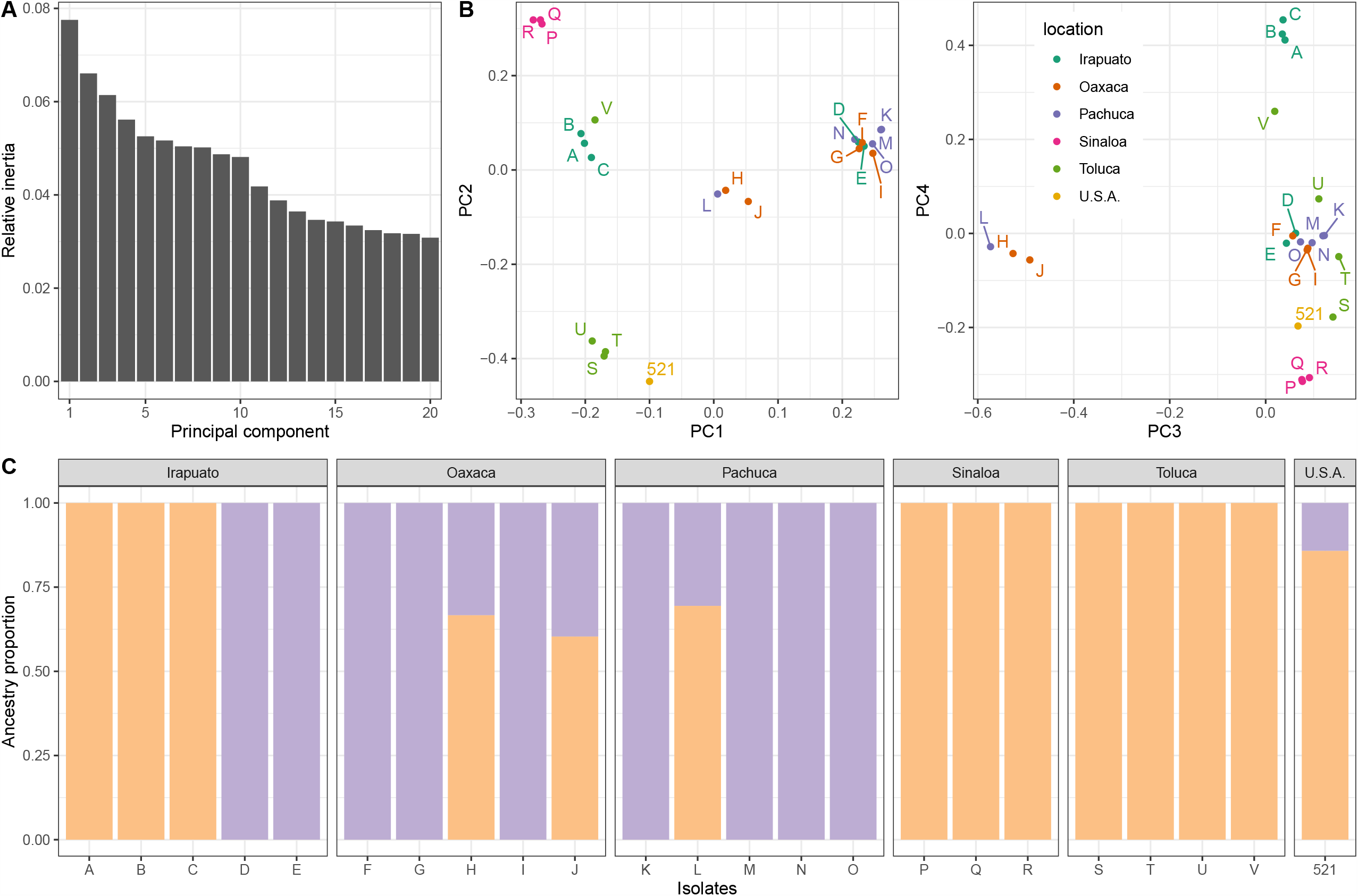
Inference of population structure reveals two subpopulations. (A) The relative inertia (vertical axis) are shown for the first 20 principal components (horizontal axis). (B) The first four principal components (PC) of a PC analysis based on genotypes are shown. The left panel shows the first and second PC, and the right panel depicts the third and fourth PC. Colors indicate the geographic origin of each isolate as shown in figure 1. (C) Population components as inferred by the ADMIXTURE model. Orange and purple colors represent the ancestry proportion of the two subpopulations (vertical axis) in each isolate (horizontal axis).

### Both subpopulations experienced strong bottlenecks

We elucidated the demographic history of the two subpopulations using a multiple sequentially Markovian coalescent (MSMC2) approach (Malaspinas *et al*., 2016), which we applied to the two population components that we inferred from the population structure analysis. We found that the ABCPQRSTUV population displayed a relatively constant effective population size between 1,000,000 and 7,000 years ago, but experienced a strong bottleneck ending about 500 years ago (figure 3A). The second subpopulation DEFGIKMNO showed a similar trend, although experiencing first a population increase before going through a stronger bottleneck starting 10,000 and ending 500 years ago. The time frame of these bottlenecks coincides with the beginning of maize domestication 6,000 to 10,000 years before present (Matsuoka *et al*., 2002; Hake and Ross-Ibarra, 2015), indicating that the demography of *U. maydis* was most likely affected by the domestication of its host plant, although to a different extent in the two populations. To assess the timing of differentiation between the two populations, we conducted a cross-coalescence analysis (figure 3B). This analysis showed that the differentiation of the two populations occurred rapidly between 5,000 and 1,000 years before present. Permutations of samples between the two populations confirmed that this pattern is very well supported by the data.

**Figure 3:**
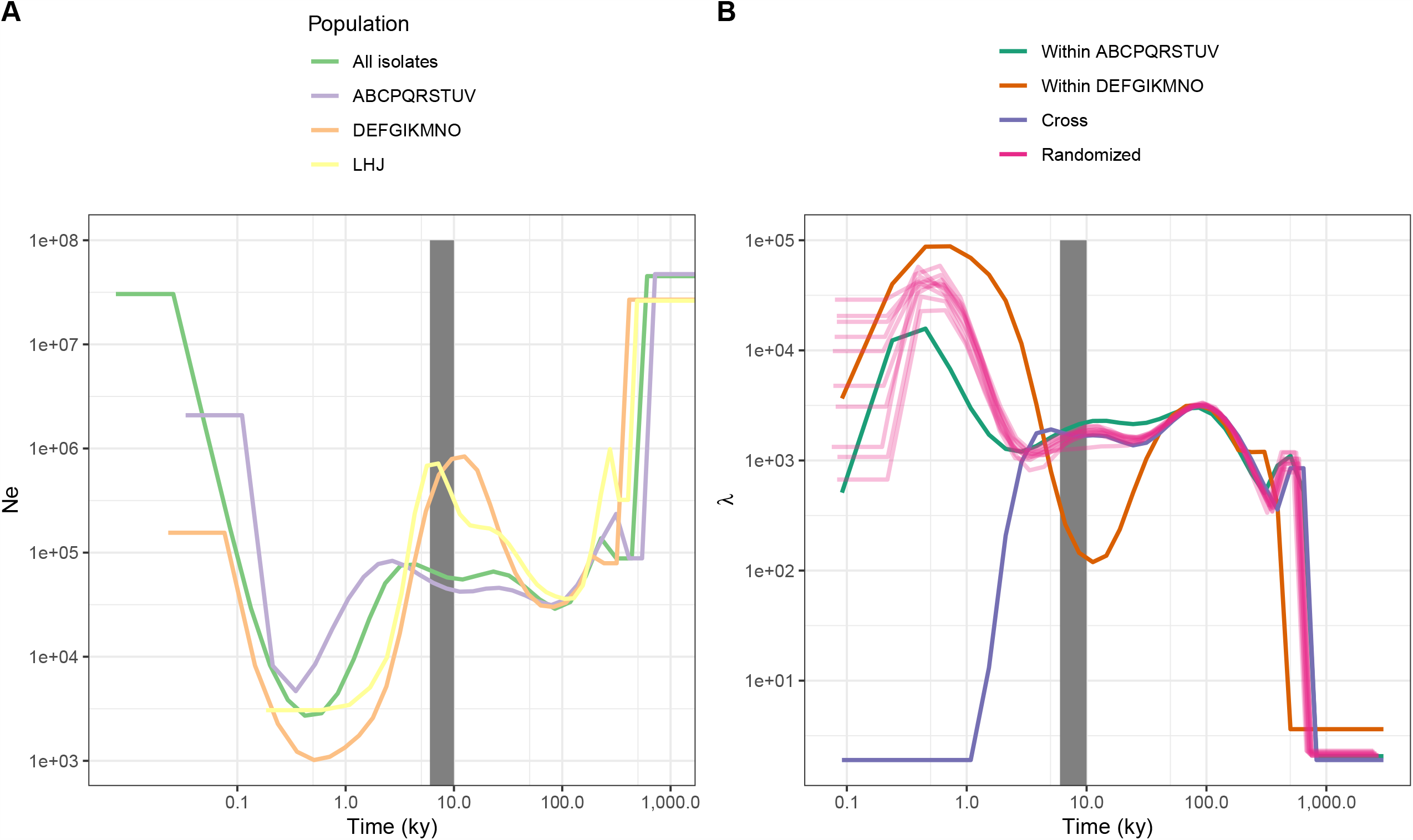
Both subpopulations experienced a recent bottleneck that started at a time overlapping with the supposed start of the domestication of maize, between 6,000 and 10,000 years ago (gray area) (A) The effective population size N_e_ (vertical axis) is shown over the last 1 million years (horizontal axis). The green line shows the result that is obtained when considering all isolates, whereas the purple, orange and yellow lines represent results for a subset of isolates as indicated. (B) A cross-coalescence analysis reveals the timing of the differentiation between the two subpopulations and shows that divergence happened rapidly within a time frame that coincides with the domestication of teosinte.

The MSMC approach infers variation of the coalescence rate in the past. Under a standard coalescent model, the coalescence rate is inversely proportional to the effective population size; thus, it provides a snapshot of the demographic history. However, assuming a standard coalescent may lead to the inference of an artificial bottleneck when the sampled population is structured (Mazet *et al*., 2016). Furthermore, the presence of loci under purifying selection can result in the inference of a recent expansion (Platt and Harris, 2020). The dataset presented here is potentially subject to these issues, given the presence of population structure and the high density of protein coding regions. While in line with the expectation that the domestication of maize impacted the evolution of the pathogen *U. maydis*, the inferred demographic scenarios should, therefore, be taken with necessary caution.

We searched for regions putatively involved in the divergence of the two populations (Wolf and Ellegren, 2017) by computing F_ST_ values in 10 kb windows along the genome (figure 4A), with the aim to highlight genes with a role in the adaptation of the pathogen during the domestication of the host. The genome-wide distribution of F_ST_ values had a mode around 0.2, and a skew toward high positive values. This distribution is well modeled by a mixture of two normal distributions (supplementary figure S1). One hundred ninety-one regions were found to be within the high-F_ST_ component with a 1% significance level, and contained 751 genes. These genes were not found to be enriched in candidate effector genes (Fisher’s exact test, *P*-value = 0.86), and GO-term enrichment analysis of these genes only exhibited top-level metabolic categories (supplementary table S7). In summary, these results provide no support for the existence of highly differentiated loci that may drive the divergence of the two subpopulations. Further analyses, notably accounting for recombination rate variation, are required to identify loci possibly involved in population differentiation (Booker *et al*., 2020).

**Figure 4:**
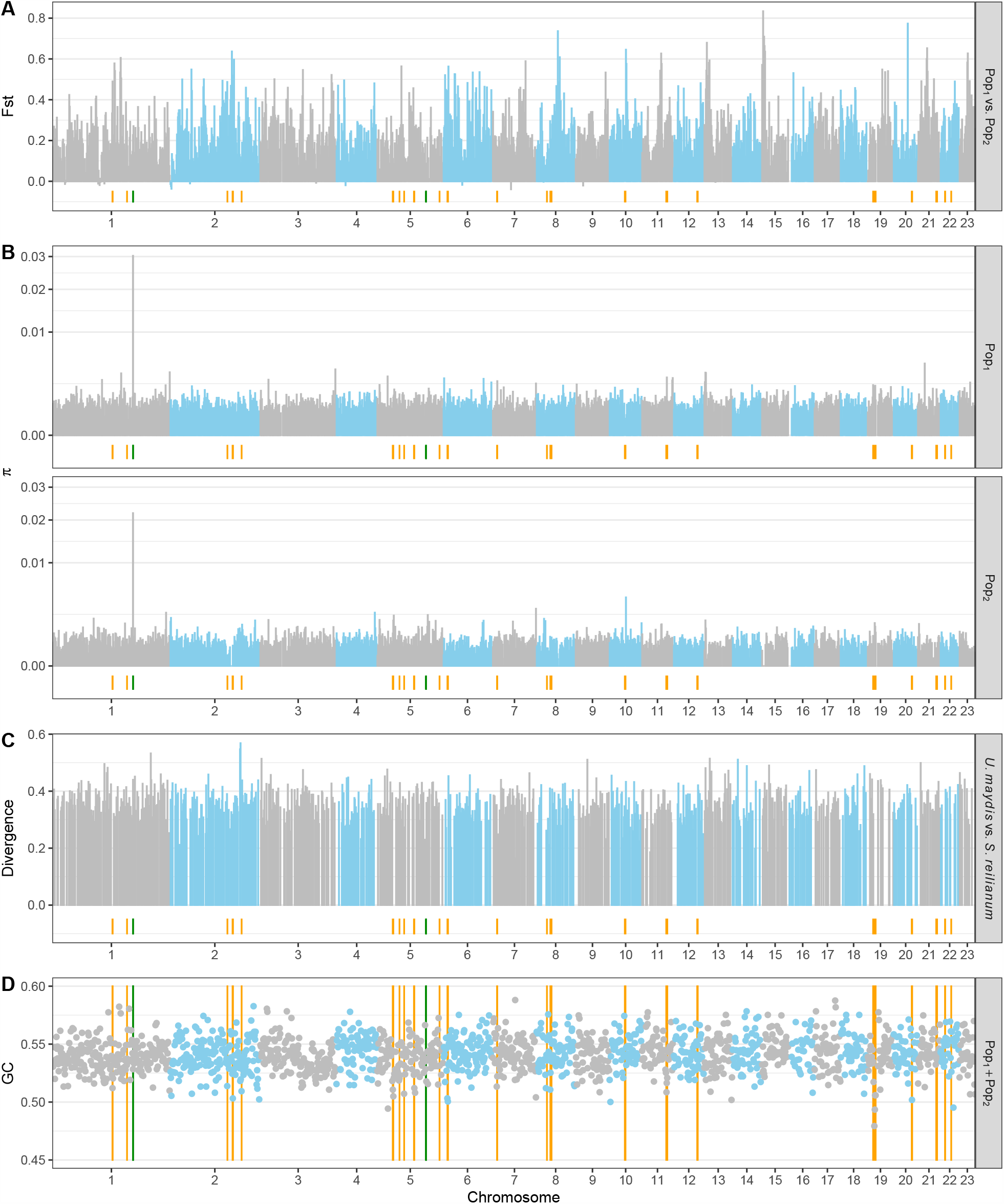
Values for different measures of genetic diversity vary at the fine (but not chromosomal) scale. The horizontal axis depicts the chromosome number, and the vertical axes show values of the fixation index (A), the nucleotide diversity in both subpopulations (B), the level of divergence between *U. maydis* and *S. reilianum* (C), and the GC content (D). Blue and gray colors of data points indicate different chromosomes. Orange lines indicate the localization of gene clusters as defined in Dutheil *et al*. (2016), and green lines show the localization of the *a* mating type locus on chromosome 5 and the *b* mating type locus on chromosome 1, respectively. Pop_1_ includes isolates A, B, C, P, Q, R, S, T, U and V and Pop_2_ the isolates D, E, F, G, I, K, M, N, and O.

### The *a* mating type locus carries alleles that are conserved within the Mexican population but different from the reference isolate

*U. maydis* possesses a tetrapolar mating system where two loci, *a* and *b*, are involved (Bakkeren *et al*., 2008). Our data set contains two idiomorphs of the *a*-locus: isolates B, D, J, K, L, M, N, O, Q, R, and U share the same *a1* idiomorph as the reference isolate, while the other isolates have another one, similar to *a2* (Bölker *et al*., 1992; Urban *et al*., 1996). Because distinct idiomorphs share no homology (Froeliger and Leong, 1991), the *a2* mating type regions could not be aligned with the reference genome and are represented as gaps in our genome alignment (supplementary file 1). The *a1* and *a2* idiomorphs, however, differ from the previously published sequences of *a1* in the 521 isolate (accession U37795) and *a2* in the RK32 isolate (accession U37796; Bölker *et al*., 1992; Urban *et al*., 1996). All Mexican *a1* alleles are 100% identical to each-other, but display a few SNPs, and a (non-coding) deletion of thymine at position 5,556 with respect to the *a1* sequence of isolate 521 (supplementary file 1). The SNPs are distributed as follows: one non-synonymous SNP is located at position eight in the *mfa1* gene (leading to asparagine instead of threonine in the a1 pheromone precursor protein). The *pra1* genes contain four SNPs in their coding regions, of which two are non-synonymous (resulting in glycine instead of alanine at position 63 and leucine instead of proline at position 103 of the amino acid sequence). The Mexican *a2* alleles differ from each other only at position 62 in the *rga2* gene, where isolates E, F, G and I encode a glycine instead of a serine. The Mexican alleles all differ, however, from the published RK32 sequence of *a2* (accession U37796). They feature a non-coding insertion of 20 nucleotides at position 1,222 and a three-nucleotide insertion at position 1,300 of the *a2* sequence from isolate RK32 (supplementary file 1), more than 2.5 kb downstream the *pra2* gene. SNPs in coding regions are distributed as follows: three in gene *pra2*, among which two non-synonymous at positions 218 (leading to alanine instead of arginine) and 294 (threonine instead of alanine), and three in *lga2*, among which one non-synonymous at position 63 (threonine instead of alanine). In addition to position 62, the *rga2* gene differs from the *a2* sequence found in isolate RK32 by two other SNPs, including a non-synonymous one at positions 45 (serine instead of isoleucine). The region between positions 1,981 to 3,735 of the U37796 sequence is notably absent in all Mexican isolates (supplementary file 1). This region contains the pseudogenized *mfa* gene copy in *a2* of RK32 reported in Urban *et al*., 1995, which was hypothesized to be a remnant of a multiallelic ancestor containing two pheromone genes. This result suggests that either the ancestor of the Mexican isolates lost this remnant, or that the degenerated pseudogenized *mfa* gene was not ancestral but inserted in the ancestor of the RK32 isolate originating from Germany.

We further observed that the assembly of the A isolate contained two complete *a1* and *a2* loci, only separated by 13 nucleotides and identical to the ones found in other Mexican isolates. We note that these 13 nucleotides include a N character, indicating that the two loci were on separate contigs and only assembled at the scaffolding stage. In order to investigate the possibility of wrong assembly or contamination, we performed a second independent DNA extraction and amplified the mating type region in isolate A, using isolate B (containing the *a1* idiomorph) as a comparison. The result revealed the presence of two amplified segments matching the sizes of the Mexican *a1* and *a2* idiomorphs, confirming their presence in the isolate, while from isolate B only one segment of the expected size for *a1* could be amplified (supplementary figure S3). This result is incompatible with the reconstructed assembly and shows that the two loci are not consecutive in the genome. To investigate possible coverage variation, we mapped all reads along the scaffold containing the *a* mating type locus in isolate A. We show that each idiomorph is supported by a coverage lower than the average of the rest of the scaffold, roughly 2/3 for the *a1* idiomorph and 1/3 for the *a2* idiomorph (supplementary figure S4). Altogether, these results suggest that the region containing the *a* mating type locus of the A isolate may have been diploid, and that the A isolate might be aneuploid. The amplified fragment corresponding to the *a2* locus is less strong than the amplified fragment representing the *a1* locus (supplementary figure 3). While this could be due to the larger size of the *a2* segment impeding the amplification efficacy, we note that it mirrors the lower coverage of the *a2* sequence in the genome sequencing (supplementary figure 4). An intriguing possibility could be that the presumed aneuploid strain is unstable and lost one of the idiomorphs during subsequent mitotic divisions. If the *a2* idiomorph is preferentially lost or if cells carrying the *a2* idiomorph are dividing more slowly than cells carrying *a1* because of autocrine pheromone stimulation, the final cell culture may contain unequal proportion of *a1* and *a2* sequences. Further genetic and cytological investigations are needed to confirm this hypothesis.

In order to assess whether the presumed diploid status of the A isolate could impact our population genomic analyses, we examined the read mapping to search for possible heterozygous positions. After mapping all reads on the assembled genome, we counted the number of mismatches in reads for each assembled position. If the genome was diploid, we would expect heterozygous positions to have alternative states in the reads, in a proportion of 50% on average. We counted the proportions of sites with an alternative state present in at least 20% of the reads: isolate A had 0.15% of such positions, while isolate B had 0.21% and isolate C had 0.20% of such sites. Scaffold 79 of strain A, which contains the *a* locus, only had 0.05% of sites with more than 20% of read supporting an alternative state. This suggests that either the potential aneuploidy of isolate A is restricted to the genomic region containing the *a* locus, or that the aneuploid region is highly homozygous. In either case, aneuploidy did not have a significant impact on the genome sequencing of isolate A outside the mating type locus.

### Genetic diversity is higher at the b mating type loci

The *b* mating type locus located on chromosome 1 represents a hotspot of diversity (figure 4C), presumably because the underlying genes evolve under balancing selection (May *et al*., 1999). Reconstructing the phylogenetic relationships between all previously sequenced alleles (described in Kämper *et al*., 2020) and alleles that could be extracted from the multiple genome alignment showed that the allele in isolate T is very similar to the *b2* allele. The *b7* allele is identified in isolates B and U. The *b14* allele is detected in isolates G, I, K, M, N, and Q. The *b15* allele is present in isolates P and R. The *b17* allele is found in isolates H, J, and L, and the *b18* allele is present in isolate V (supplementary figure S5). Interestingly, two alleles, one from isolates A and C and one from isolates D, E, F, O, and S did not cluster with any previously identified allele and may therefore be potentially novel. This finding needs to be corroborated with *in vitro* mating assays and successful plant infections.

### The GC content of U. maydis correlates with chromosome size and is lower in virulence clusters

We investigated several patterns of genetic diversity in windows of 10 kb along the genome of *U. maydis*. The GC content, divergence with *S. reilianum* and mean number of nucleotide differences within the two subpopulations all appeared to be homogeneous at the chromosome scale (figures 4B-D). We report a slightly significant negative correlation between the average GC content per chromosome and the chromosome length (Kendall’s rank correlation test, tau = −0.296, *P*-value = 0.04984). A negative correlation between chromosome size and average recombination rate is frequently observed in eukaryotes and usually explained by small chromosomes having a higher recombination rate (Kong *et al*., 2002; Jensen-Seaman *et al*., 2004). A positive correlation between GC content and recombination rate could be the result of GC-biased gene conversion (Lesecque *et al*., 2013), and this could drive the higher GC content in smaller chromosomes. Finally, we report that the GC-content is significantly lower in regions encompassing clusters of effectors (Wilcoxon’s rank test, *P*-value = 1.372×10^−14^), in agreement with the previous report that virulence clusters are associated with AT-rich repeat elements (Dutheil *et al*., 2016). Below, we further discuss the link between repeat elements and virulence clusters.

### Genetic diversity is higher around clusters of effector genes

In *U. maydis* and related species, several effector genes are organized in gene clusters, and many of them have a role in virulence (Kämper *et al*., 2006; Schirawski *et al*., 2010). We previously reported that, when comparing multiple related species, these gene clusters are enriched in genes evolving under positive selection (Schweizer *et al*., 2018). Here, we find that the mean number of nucleotide differences was on average higher in windows overlapping with a virulence cluster: the median across 10 kb windows of the mean number of nucleotide differences is 0.0011 in virulence cluster regions, compared to 0.0008 for the rest of the genome in population ABCPQRSTUV (Wilcoxon test, *P-*value = 5.131×10^−7^), and 0.0007 vs. 0.0004 in population DEFGIKMNO (Wilcoxon test, *P-*value = 0.02154), consistent with a previous report based on a smaller set of genes (Kellner *et al*., 2014). This higher diversity may result from a locally higher mutation rate and / or distinct selection regime. A higher mutation rate can be caused by the activity of transposable elements (TEs), directly or indirectly: active TEs may locally introduce mutations, the generated occurrence of repeated sequence may lead to increased slippage of the DNA polymerase, and mechanisms protecting the genome against TEs may be leaky and affect surrounding regions (Horns *et al*., 2012; Laurie *et al*., 2012; Dutheil *et al*., 2016). In addition, selection may impact diversity along the genome (Charlesworth, 2009; Gossmann *et al*., 2011; Grandaubert *et al*., 2019). All these mechanisms may act simultaneously and are difficult to disentangle. Additional insights may be obtained by contrasting different classes of polymorphisms in protein coding regions.

### Clustered effector genes show a high rate of adaptive mutations

To elucidate the selection regime under which different categories of genes evolve, we assigned each gene in *U. maydis* to one of the three categories “clustered genes” (genes in predicted virulence clusters), “non-clustered effectors” (genes predicted to encode an effector protein, but not located in a predicted cluster) or “remaining genome” as defined earlier (Dutheil *et al*., 2016). We then computed the amount of synonymous and non-synonymous divergence, which we compared to the amount of synonymous and non-synonymous polymorphism for each category of genes. We find no significant difference of synonymous nucleotide diversity (π_S_, figure 5A), non-synonymous diversity (π_N_, figure 5B), and ratio of non-synonymous to synonymous diversity (π_N_/π_S_, figure 5C) between genes in the three categories. This contrasts with results obtained in other fungal pathogens such as *Z. tritici*, where genes encoding effector proteins are evolving under relaxed purifying selection evidenced by a higher π_N_/π_S_ ratio (Grandaubert *et al*., 2019). The ratio of non-synonymous to synonymous divergences (d_N_/d_S_ = ω), however, was found to be significantly higher in clustered genes than in non-clustered effectors, which have a significantly higher d_N_/d_S_ than non-effector genes (figure 5D; *P*-value = 0.00599). In order to disentangle the adaptive and non-adaptive part of the d_N_/d_S_ ratio, we fitted models of distribution of fitness effects on polymorphic data for genes in each category (Galtier, 2016), and estimated the rate of non-adaptive non-synonymous substitutions (ω_NA_), the rate of adaptive non-synonymous substitutions (ω_A_), and the proportion of non-synonymous substitutions that are adaptive (α). We find that genes located in clusters of effectors have a lower rate of non-adaptive non-synonymous mutations (model averaged ω_NA_ = 0.06, Figure 5E and supplementary table S6) compared to non-clustered effector genes (ω_NA_ = 0.19; *P*-value = 0.042) and non-clustered non-effector genes (ω_NA_ = 0.12). The rate of adaptive mutations was found to be significantly higher in genes located in effector clusters (ω_A_ = 0.38, Figure 5F), compared to non-clustered effectors (ω_A_ = 0.11; *P*-value = 0.00599) and non-clustered, non-effector genes (ω_A_ = 0.11; *P*-value = 0.00599). The proportion of adaptive non-synonymous substitutions (α = ω_A_ / ω) was found to differ marginally between genes in effector clusters (α = 86%, Figure 5G), non-clustered effector genes (α = 46%; *P*-value = 0.0599) and non-clustered, non-effector genes (α = 48%; *P*-value = 0.0879). These results are consistent with previous reports that genes encoding effector proteins undergo a higher rate of adaptive evolution (Stukenbrock and Bataillon, 2012; Grandaubert *et al*., 2019). Differences in evolutionary rates can be explained by differences in the effective population size, *Ne*, or by the selection coefficient, *s*, of the mutations. Consequently, two non-exclusive hypotheses for this observation can be invoked. First, because some effector genes are essential for virulence, the fitness effect of mutations at these genes might be larger. Under an arms race scenario, mutations creating new alleles that prevent recognition by the host will have a large positive effect (large positive value of *s*, leading to a high value of ω_A_). When established, however, successful effector alleles are under strong negative selection so that most mutations are highly deleterious (larger negative value of *s*, impeding the accumulation of non-adaptive mutations and leading to a low value of ω_NA_), until the plant target protein evolves a sequence that prevents interaction with the effector. Examples for such conserved effectors include Pep1 (Doehlemann *et al*., 2009) and Pit2 (Doehlemann *et al*., 2011), which are inhibitors of the maize peroxidase POX12 and a group of papain-like cysteine proteases, respectively (Hemetsberger *et al*., 2012; Müller *et al*., 2013; Misas Villamil *et al*., 2019). Other examples of important effectors are ApB73 and Cce1, but their molecular function remains to be elucidated (Stirnberg and Djamei, 2016; Seitner *et al*., 2018). Second, the pattern of selection may result from the genomic context of the underlying genes. A higher recombination rate in the region of effector clusters is predicted to locally increase *Ne* by reducing linkage disequilibrium and, therefore, to increase the efficacy of both negative and positive selection (Charlesworth and Campos, 2014; Croll *et al*., 2015; Grandaubert *et al*., 2019). However, we could not infer a recombination map based on the 22 Mexican *U. maydis* isolates, because the small sample size and low genetic diversity prevented the application of linkage disequilibrium-based methods (Baudat *et al*., 2013). Moreover, the small chromosome and genome size as well as the high density of protein coding genes (about 61.2 % of the genome) did not allow us to obtain reliable results with approaches based on sequentially Markovian coalescent processes like iSMC (Barroso *et al*., 2019). Constructing a recombination map for *U. maydis*, therefore, would require the sampling of more isolates from diverse geographic origins.

**Figure 5:**
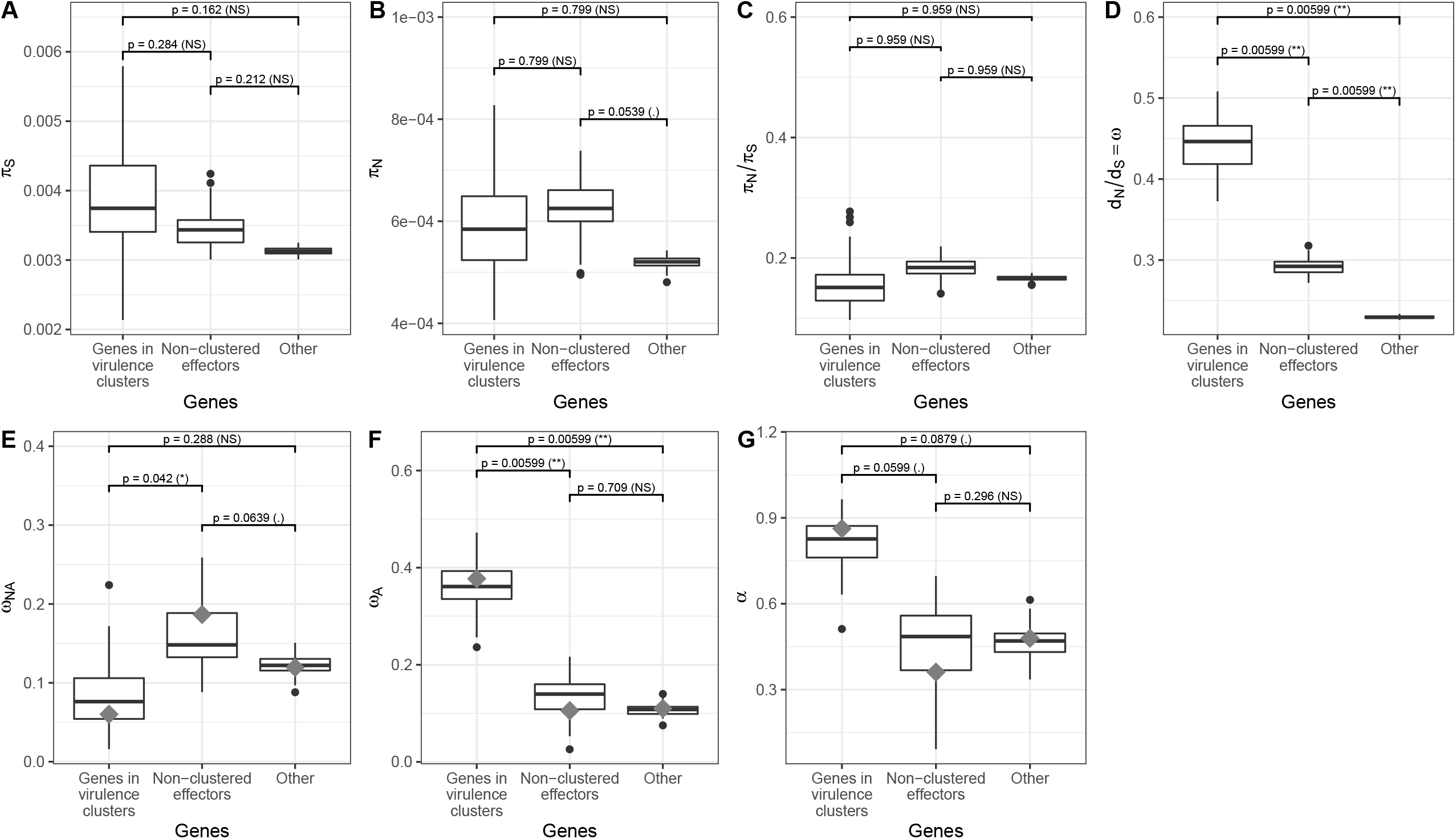
Genes in virulence clusters display a higher rate of adaptive substitutions. *P*-values for pairwise comparisons between clusters of effector genes, unclustered effector genes and non-clustered, non-effector genes are computed as described in Methods. (A) Mean number of synonymous nucleotide differences (π_S_), (B) mean number of non-synonymous nucleotide differences (π_N_), (C) ratio of the mean number of non-synonymous nucleotide differences and mean number of synonymous nucleotide differences (π_N_/ π_S_), (D) ratio of non-synonymous over synonymous divergence (d_N_ / d_S_), (E) rate of non-adaptive non-synonymous substitutions (ω_NA_), (F) rate of adaptive non-synonymous substitutions (ω_A_), (G) proportion of non-synonymous adaptive substitutions (α). Box-and-whiskers plots indicate the median, first and third quantiles over 100 bootstrap replicates performed using the best model according to Akaike’s information criterion. Grey diamonds indicate the corresponding value when averaging over all models of distribution of fitness effects (see Methods for details).

Effector genes are characterized by their relatively short size, and genes in virulence clusters are, therefore, significantly shorter than non-clustered genes (317 amino-acids on average *vs*. 396 amino-acids for non-clustered genes, Wilcoxon test, *P*-value < 2.2×10^−16^). Short genes have been shown to evolve faster, as the encoded proteins have a higher proportion of exposed residues, which tend to experience a higher frequency of adaptive substitutions (Moutinho *et al*., 2019). In order to test whether the shorter protein size of genes in virulence clusters explains their higher rate of adaptive evolution, we created samples of genes with similar lengths to that of genes in virulence clusters (see Methods and supplementary file S1). We then conducted the same analyses on the size-restricted data sets. The results, including estimates of ω_A_ and ω_NA_ are very similar when controlling for protein length, suggesting that it does not account for the observed differences (supplementary figure S6).

Lastly, we note that predicted effector genes that are not located in virulence clusters do not show an increased rate of adaptive evolution, contrasting with other pathogens where effector-encoding genes show an increased evolutionary rate (Stukenbrock *et al*., 2011; Hacquard *et al*., 2012; Huang *et al*., 2014; Sperschneider *et al*., 2014; Sharma *et al*., 2014). The *U. maydis* genome is almost devoid of transposable elements (TEs), which are mostly concentrated in the region of virulence clusters. This distribution may result from TEs being locally tolerated, as they could be providing a supply of mutations in rapidly evolving regions. Another explanation is that negative selection against TEs is locally not strong enough, owing to recurrent positive selection on genes in virulence clusters, which reduces the local effective population size because of linkage. These two explanations are not exclusive and can both contribute positively to the local accumulation of TEs. Altogether, these results highlight the singular role that virulence clusters play in the adaptation of *U. maydis* to its host.

## Acknowledgements

This work was generously supported by funds from the Max Planck Society. We thank Octavio Paredes-López for providing the Mexican isolates of *U. maydis*.

## Author contributions

Conceptualization: GS (equal), JYD (equal), RK (equal); Formal Analysis: GS (equal), JYD (equal); Funding Acquisition: RK (lead); Investigation: GS (equal), JYD (equal), MBH (supporting), GVB (supporting), NR (supporting), KM (supporting), RK (supporting); Project Administration: GS (lead), JYD (supporting), RK (supporting); Supervision: JYD (lead), RK (supporting); Visualization: JYD (lead), GS (supporting); Writing – Original Draft Preparation: GS (lead), JYD (supporting), NR (supporting), KM (supporting), MBH (supporting); Writing – Review and Editing: GS (equal), JYD (equal), RK (equal), GVB (supporting)

## Data availability

Illumina paired-end reads and assembled genome sequences were deposited at NCBI (BioProject Id: PRJNA561077). The Whole Genome Shotgun project has been deposited at DDBJ/ENA/GenBank under the accessions WEIZ00000000 to WEJU00000000. The versions described in this paper are versions WEIZ01000000 to WEJU01000000. Multiple genome alignments, sequences of open reading frames that could be extracted from the multiple genome alignment, and alignments of *U. maydis* gene sequences with their outgroup in *S. reilianum* are available from supplementary file 1. Scripts necessary to reproduce the analyses are available at http://gitlab.gwdg.de/molsysevol/UmaydisPopGen and in supplementary file 1.

## Supplementary figures

Supplementary figure S1: Distribution of F_ST_ values in 10 kb windows. The orange curve shows the fit of a two-normal mixture distribution.

Supplementary figure S2: Detailed results from the ADMIXTURE analysis, as represented by the PONG software. Each row represents the most likely population structure for a given number of subpopulations (K). Model fitting was repeated 10 times in each case with different initial random parameter values, and the most frequently inferred scenario was depicted. Ratios in blue indicate the number of replicates supporting the represented scenario in each case.

Supplementary figure S3: Amplification of the *a* mating type locus. (A) Expected sizes in the reference isolates for the two idiomorphs *a1* and *a2*, the Mexican isolates B and C (having the Mexican *a1* and *a2* idiomorphs) and isolate A, predicted to carry both idiomorphs. Expected sizes of the Mexican isolates are based on the genome assemblies. (B) Amplification of the corresponding locus in isolates A (lane 1) and B (lane 2). M, size marker.

Supplementary figure S4: Sequencing coverage along scaffold 79 of the A isolate assembly. The region containing the *a1* idiomorph is depicted in red, the region containing the *a2* locus is plotted in green, and the central region of the scaffold (N) in blue. Coverage is measured in number of reads mapped to each position. (A) Per nucleotide coverage. Straight lines represent the linear regression on the corresponding data points. (B) Coverage distribution per region.

Supplementary figure S5: Phylogenetic tree of the mating type region of the b-locus. Left tree: *bW* alleles (*UMAG_00578*). Right tree: *bE* alleles (*UMAG_12052*). Nodes with a bootstrap value lower than 60% have been unresolved and displayed as multifurcations. Colors of the isolate names distinguish the ABCPQRSTUV and DEFGIKMNO subpopulations, as well as the three admixed individuals H, J, and L. Alleles of the reference strain 521 are is displayed in black, and all previously published alleles are in grey boxes.

Supplementary figure S6: Comparison of genetic diversity and rate of adaptive substitutions in clusters of effector genes, unclustered effectors and non-clustered, non-effector genes after controlling for protein length. Legend as in Figure 5.

## Supplementary tables

Supplementary table S1: Overview of the 22 Mexican *U. maydis* isolates

Supplementary table S2: Overview of the assembly results with different kmer lengths

Supplementary table S3: Intermediate alignment lengths and number of blocks

Supplementary table S4: List of all *U. maydis* genes with mapped Interpro domains, Gene Ontology Terms, F_ST_ values, and outgroup sequences

Supplementary table S5: SiLiX results for different identity and coverage settings

Supplementary table S6: Estimates of the rate of adaptive substitutions for various models of distributions of fitness effects.

Supplementary table S7: Gene Ontology Term enrichment analysis for genes with high F_ST_ values

## Supplementary files

Supplementary file S1: Data set and scripts necessary to reproduce the analyses in this work.

## Notes

### Competing Interest Statement

The authors have declared no competing interest.

