## Supplementary figures and images for "Population genomics of the maize pathogen *Ustilago maydis*: demographic history and role of virulence clusters in adaptation"

### Supplementary Figure S1

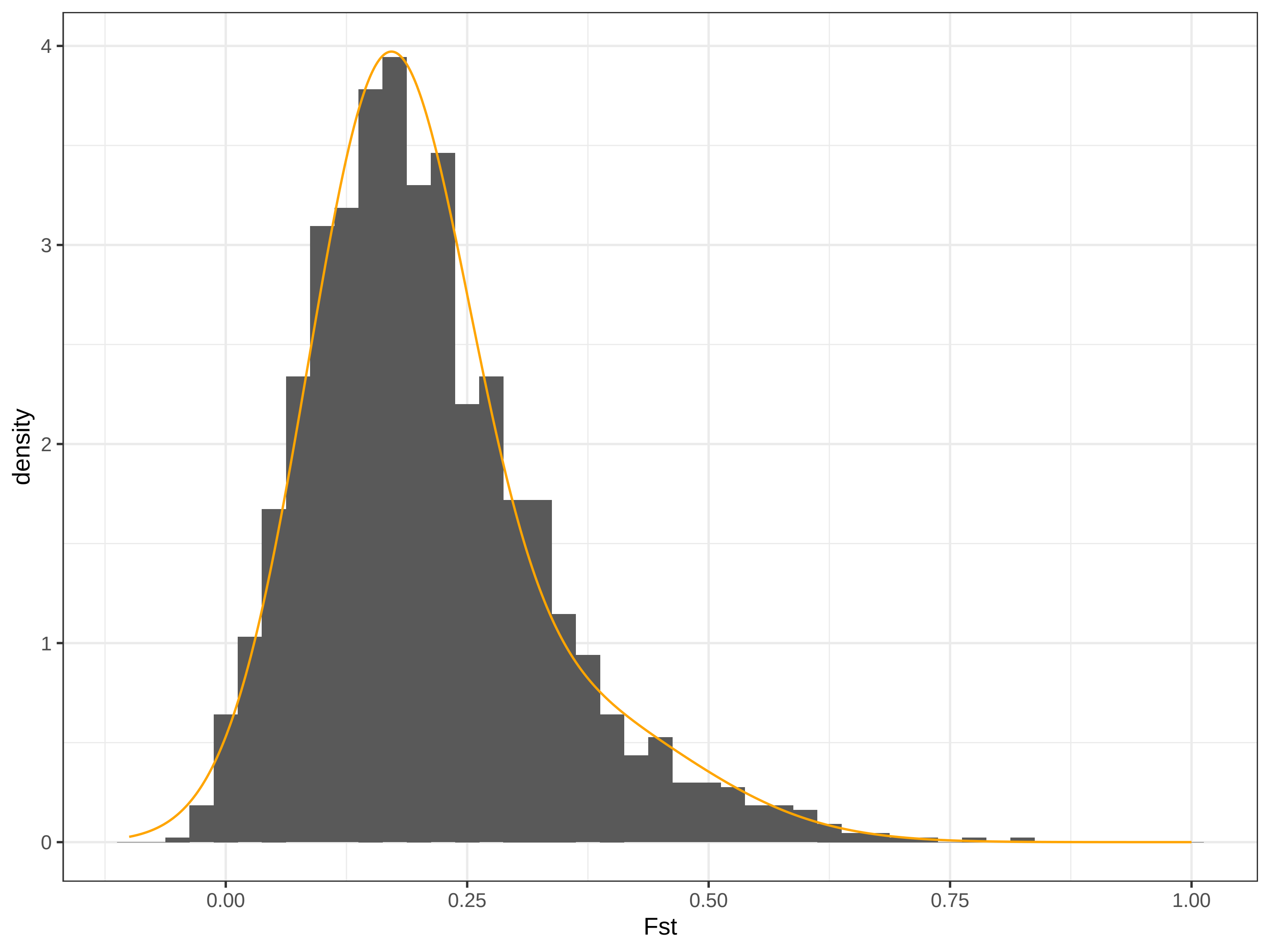

### Supplementary Figure S2

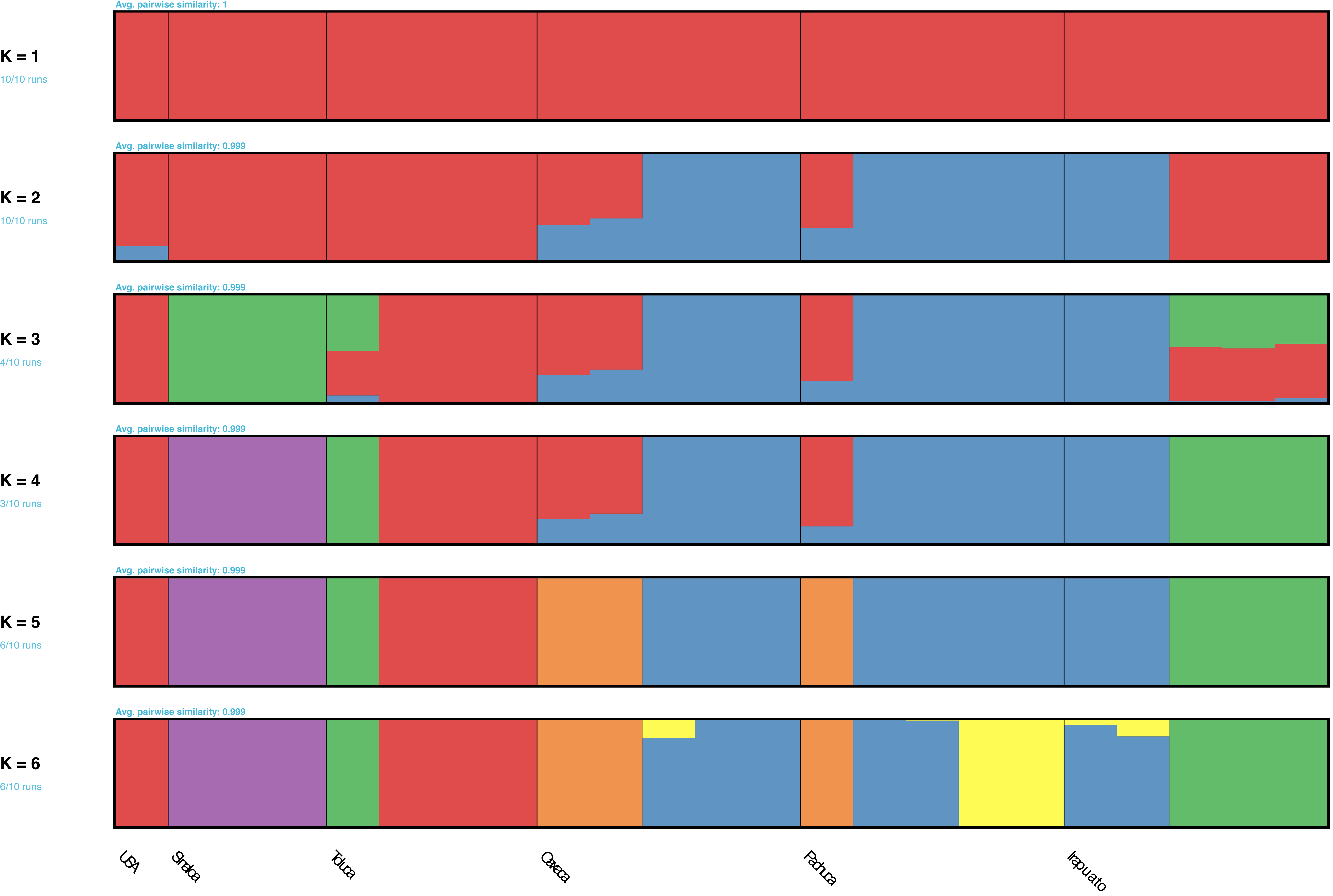

### Supplementary Figure S3

**A**

| Strain | Expected size |
|---|---|
| Reference a1 | 4,961 bp |
| A (a1 + a2) | 11,890 bp |
| B (Mexican a1) | 4,959 bp |
| Reference a2 | 8,809 bp |
| Mexican a2 | 7,082 bp |

**B)**

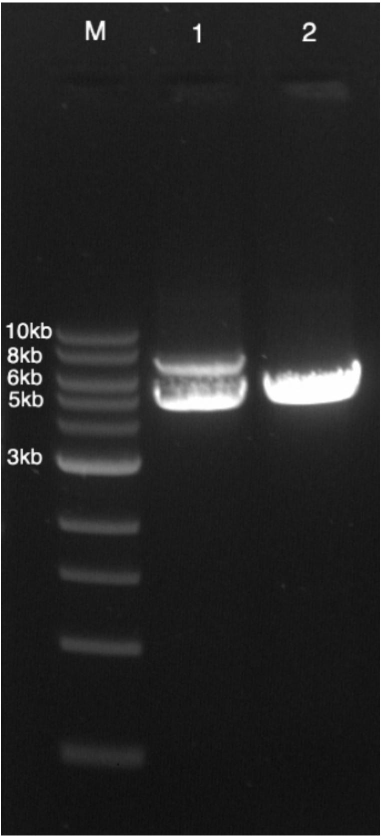

### Supplementary Figure S4

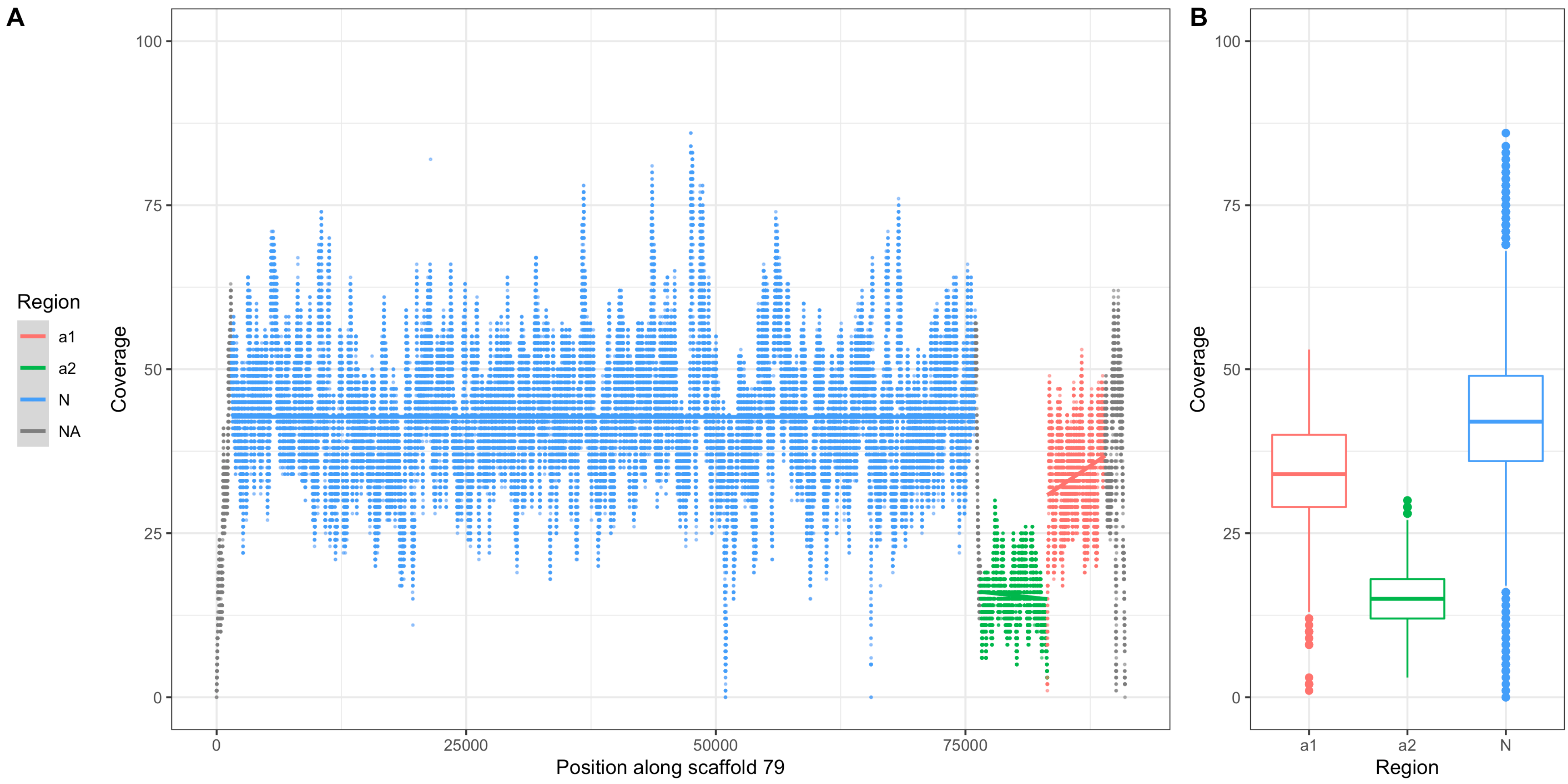

### Supplementary Figure S5

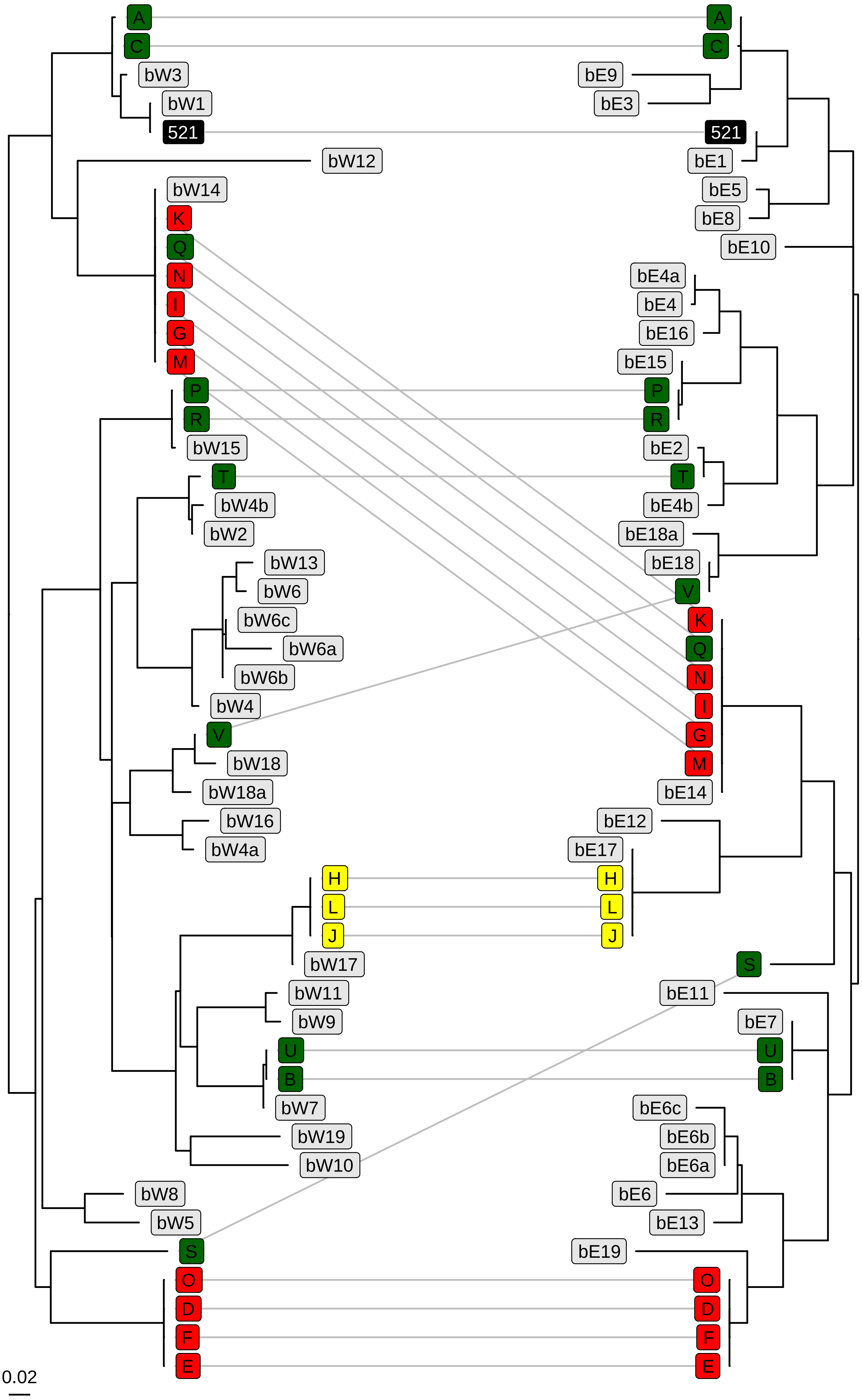

### Supplementary Figure S6

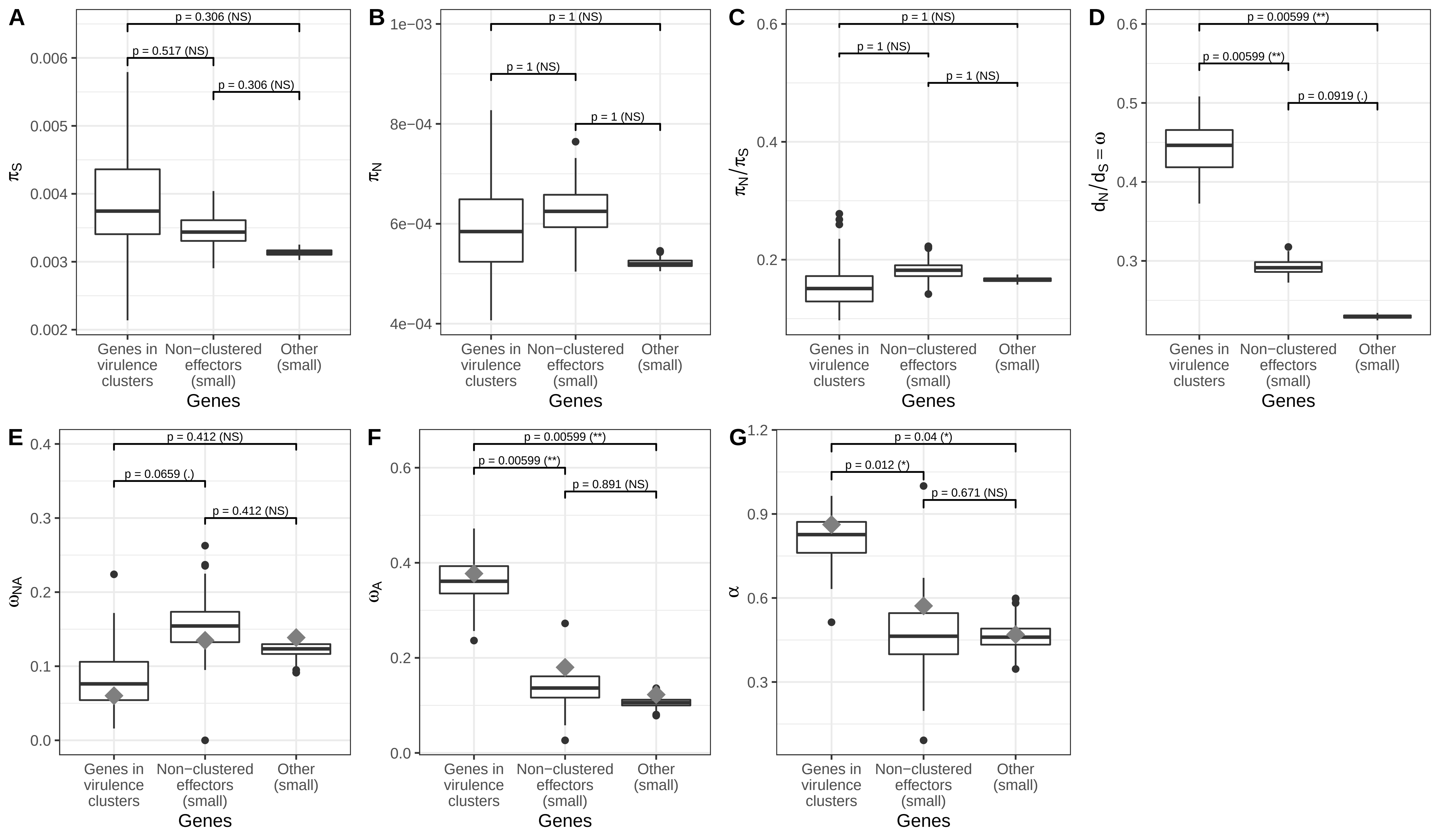
